# Analysis of the Functional Differentiation Sites in Vertebrate Neuronal Nicotinic Acetylcholine Receptor Subunits

**DOI:** 10.1101/2020.10.22.350595

**Authors:** Mengwen Zhao, Yuequn Ma, Juncai Xin, Changying Cao, Ju Wang

## Abstract

The nicotinic acetylcholine receptors (nAChR) belong to a large family of ligand-gated ion channels and are involved in the mediation of fast synaptic transmission. Each receptor is made up of five subunits that arrange symmetrically around a central pore. Despite the similarity in their sequences and structures, the properties of these subunits vary significantly. Thus, identifying the function-related sites specific to each subunit is essential for understanding the characteristics of the subunits and the receptors assembled by them. In this study, we examined the sequence features of the nine neuronal nAChRs subunits from twelve representative vertebrate species. Analysis revealed that all the subunits were subject to strong purifying selection in evolution, and each was under a unique pattern of selection pressures. At the same time, the functional constraints were not uniform within each subunit, with different domains in the molecule being subject to different selection pressures. Via evolutionary analyses, we also detected potential positive selection events in the subunits or subunit clusters, and identified the sites might be associated with the function specificity of each subunit. Furthermore, positive selection at some domains might contributed to the diversity of subunit function; for example, the β9 strand might be related to the agonist specificity of α subunit in heteromeric receptor and β4-β5 linker could be involved in Ca^2+^ permeability. Subunits α7, α4 and β2 subunits possess a strong adaptability in vertebrates. Our results highlighted the importance of tracking functional differentiation in protein sequence underlying functional properties of nAChRs. In summary, our work may provide clues on understanding the diversity and the function specificity of the nAChR subunits, as well as the receptors co-assembled by them.

## 1. Introduction

Nicotinic acetylcholine receptors (nAChRs) are ligand-gated ion channels responsible for conducting neurotransmission activated by ligands like acetylcholine or nicotine [1–4]. By translating the binding of ligands into a flow of certain ions across the membrane and inducing a specific cellular response, these receptors play fundamental roles in a diversity of physiological processes. They are also involved in various neuropathic diseases such as Alzheimer’s disease, Parkinson’s disease, schizophrenia, autism and nicotine addiction [5–8], and are suggested to be potential targets of therapeutic agents [9]. Altogether, seventeen nAChR subunits have been identified in vertebrate, among which the α2-α7 and β2-β4 subunits, usually referred to as the neuronal nAChR subunits, are widely expressed throughout the brains. [10]. These subunits usually co-assemble into homo- or hetero-pentameric receptors, like the α7* and α4β2* (* denotes any other subunits) subtypes, which results in a large number of possible combinations.

All the subunits share a similar molecular structure. Generally, the framework is composed of an amino-terminal extracellular domain (ECD), four transmembrane domains (termed as TM1-TM4), a large and variable intracellular domain (ICD) between TM3 and TM4, as well as a short extracellular carboxyl-terminus. The TM2 helices of the five subunits in a receptor are closely packed to pave the inner circle of the ion channel pore, while TM1, TM3 and TM4 concentrically form the outer circle [11–15]. The intracellular domain, together with some parts of the extracellular domain, may impart features specific to each subunit [16–18].

Despite the similarity in their sequences and structures, properties of these subunits vary greatly. Some subunits (e.g., α2-α4, α6 and α7) can serve as the principal components harboring the ligand-binding sites. Some subunits (e.g., α5 and β3) are structural subunits that do not participate in ligand binding; instead, they impart unique functional and pharmacological properties to the receptors [16, 19–20]. While subunits such as α7 are able to form homomeric receptors, most other subunits can only co-assemble with other types of subunit to generate functional receptors. In addition, subunits contribute to the variability of the receptors in many aspects. For example, receptors containing subunit α4 have a high affinity to nicotine [21–22], and the homomeric receptor formed by α7 displays a marked permeability to calcium ions and is highly sensitive to the antagonist α-bungarotoxin [23–25].

Thus, the assembly of neuronal nicotinic receptors and their molecular kinetics properties, ion channel permeability, as well as pharmacological specificities, are closely related to the common and specific features of the subunits [26]. Identifying the molecular features specific to these structurally homologous and topographically similar nAChR subunits, will not only help us understand how the subunits are assembled into diverse receptors of variable properties, but also provide useful information on how to design more effective therapeutic agents.

However, our understanding on the correlation between the subunit features and the diversity of their property and function, as well as the subunit combination in nAChRs, is still far from complete. Available evidence indicates that all the current nAChR subunits evolved from a common ancestor, which appeared first in the nervous system and functioned as a homo-oligomer similar to the present homo-pentameric receptors formed by subunit α7 [12]. Genes encoding nAChR subunits might be generated or disappeared in evolution, eventually forming the diversification of the subunits and holding the main sequence identity [27–28]. The same process also confers unique subunit composition, specialized properties and functions to each type of receptor.

With the availability of more and more data on the sequence and structure of nAChR subunits, it becomes feasible to investigate the correlation between the sequence characteristics and functional specificity of the subunits by analyzing their molecular evolution features [12–13, 29–32]. While most of the available phylogenetic analyses focus on the origin, duplication, gene structure of nAChR subunits, as well as their evolutionary conservation, divergence and regulation, few studies have been conducted to analyze the specific features of the subunits. Tsunoyama and Gojobori proposed that the evolutionary pressure imposed on the nAChR subunits varied with their relative importance in function [13]. By comparing the ratio of the number of nonsynonymous substitutions to that of synonymous substitutions between sequences of subunits, they found that in the muscle system, subunit α1 and ε were more conservative than others, consistent with their roles in ligand-binding; while in the nervous system, subunit α7 was subject to the strongest functional constraint [13].

Since the global functions of nAChRs, such as subunit cooperativity and compatibility, likely emerged from a network of amino acid residues distributed across the entire pentameric complex, a comparative approach that is able to trace the evolutionary history of the subunits and allows for rational mapping of biological relevant features within the sequence space, may offer new hope for uncovering the amino acid origins of these enigmatic properties [33].

In the present work, by analyzing the protein sequences of neuronal nAChR subunits expressed in human and other vertebrate brain, we detected functional constraints in different domains of each subunit, and provided quantitative evidence on their functional differences. In addition, we also identified a site profile that may be responsible for the functional differentiation among these subunits.

## 2. Materials and Methods

### 2.1. Sequences collection and phylogenetic analysis

We analyzed the neuronal nAChR subunits α2-α7 and β2-β4 from 12 representative vertebrate species, i.e., *H.sapiens* (Hs), *G.gorilla* (Gg), *M.mulatta* (Mam), *R.norvegicus* (Rn), *M.musculus* (Mum), *C.familiaris* (Cf), *M.domestica* (Md), *G.gallus* (Gag), *A.mississippiensis* (Am), *X.laevis* (Xl), *L.chalumnae* (Lc), and *D.rerio* (Dr). These species are common model organisms, and a full range of neuronal nAChRs genes have been found in their genomes. For these species, we first retrieved the protein coding sequences of the nine neuronal nAChR subunits identified in human from the NCBI Nucleotide database (https://www.ncbi.nlm.nih.gov/nucleotide/), and then used them as templates to query the corresponding homologous sequences in the other species. For subunit α4 of *M.mulatta*, α6 of *L.chalumnae* and α7 of *G.gorilla*, the nucleotide sequences were unavailable. For each subunit, the peptide sequence corresponding to the nucleotide sequence was also retrieved. Thus, a total of 105 pair of nucleotide/protein sequences were collected (Supplemental Table 1). Multiple sequence alignments (MSA) were performed on the peptide sequences using ClustalW by aligning codon included in MEGA 7.0 package with default parameters [34]. Maximum Likelihood (ML) method was used to infer phylogenetic tree and bootstrap analysis was conducted with 1000 replicates to verify the significance of nodes (Supplementary Figure S1 and S2). To provide a 3D view of the protein structure, we employed the X-ray structure of the human α4β2 nicotinic receptor acquired from the Protein Data Bank (PDB) (5KXI) (Supplementary Figure S3) [35].

### 2.2. Branch-site analysis of positive selection

We calculated the ratio of non-synonymous to synonymous substitution rate (ω=dN/dS) [36], a measure of selective pressure, for each of the nine subunits based on the codon-substitution model and maximum likelihood analysis via the CodeML method of PAML software [37]. Under the framework of Markov process, calculation of ω is based on the transition rate of codon *i* to codon *j*, weighted by the prior probability of observing codon *j*. For each codon, there are 61 possible transition rates, so all the rates can be laid out in a 61×61 transition rate matrix Q [36]. For each subunit, every amino acid in the sequence is assigned an ω value; and an ω value significantly higher than 1 is viewed as evidence for positive selection, ω < 1 suggests purifying selection and ω =1 means neutral evolution.

According to the branch-site model proposed by Yang et al. [38], the ω values is allowed to vary both among sites (codons in the sequence) in a protein and across the branches on the tree, as long as the foreground (the branch of interest) and background (all other branches) lineages are specified *a priori* within the phylogeny. In the analysis, each site in the protein is assigned to one of four classes. Sites Class 0 and 1 are codons conserved (< ω < 1) or neutral (ω = 1) throughout the tree and have the same ω value in the background and foreground lineages. Site Class 2a and 2b include codons that are conserved or neutral on the background branches, but are under positive selection on the foreground branches with ω > 1. To test the significance, positive selection is allowed on the foreground branches and is compared with a null distribution in which ω is set to 1 to get rid of the effect of neutral selection [38].

Log-likelihood of both models act as criterion. P-value was assessed for the λ^2^ distribution of LRT (twice the log-likelihood difference between the test model and the null model) with the degree of freedom equal to the difference in parameters between the compared two models [39].

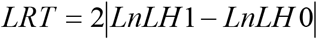

### 2.3. Sites models analysis

The site model was adopted to examine the positive selection among sites. Three pairs of models were compared, i.e., M0-M3, M1a-M2a and M7-M8. First, a single ω value was obtained for the whole phylogenetic tree at all sites using the M0 model. Then M0 was compared with M3 model to test the variation of ω. The M1a model assumes two site classes, ω <1 and fixed ω=1, as well as the proportions under two site classes. The M2a model is an extension of M1a model, with a third site class of ω >1 added. The M7 assumes these sites is in accordance with a beta distribution under neutral or purifying selection cases, and M8 adds a class of sites allowing positive selection from the real data [40]. Three pairs are compared by means of the LRT to accept or reject the hypothesis of the null model. The ω value and LRT express the sign of the adaptive selective pressure and the probability that accounts for functional site or functional constraint.

## 3. Results

### 3.1. Evaluation of functional constraint on neuronal nAChR subunits

Consistent with previous findings, our analysis indicated that the sequences of the neuronal nAChR subunits were quite conserved in evolution. Especially, the transmembrane domains were remarkably conserved, with a large fraction of non-polar residues (e.g., Phe, Val, Ile and Leu). The coupling region, i.e., the interface between the ECD and the TMDs, were also conserved across all the subunits (segments shown red in Supplemental Figure S1). Some conserved specific-subunit motifs were also detected in ICD, such as ‘LFM’ and ‘RRQR’ of β2, ‘RSLSVQ’ of α4, and ‘RSSSSES’ of α3 (segments shown in green in Supplemental Figure S1).

To obtain a better understanding on the evolution mode and their roles of the neuronal subunits, we evaluated the functional constraints on each subunit by analyzing the selective pressure imposed on it in evolution, which was measured by the ratio of non-synonymous to synonymous substitution rate ω. We obtained the ω values for each site along the sequence of every subunit across the 12 species. Then, a window-based analysis of ω values was performed to measure the local variation of selection pressure, with the window size defined as a sequence segment of 15-codon on the alignment of the same type of subunit (Figure 1). For each subunit, a clear distribution pattern of ω values corresponding to the ECD, TM1-TM3, ICD, TM4, as well as the extracellular C-terminus, was revealed. Generally, the ω values of all the segments were significantly smaller than 1.0, indicating that the presence of strong global purifying selection on all the subunits. However, the functional constraint on each subunit was not uniform. The ω values corresponding to transmembrane domains TM1, TM2 and TM3 of each subunit were close to zero and approximately parallel to the X-axis, indicating these domains were quite conserved during evolution. On the other hand, although the TM4 domain of most subunits was also conserved, subunits such as α6, β3 and β4, had obviously larger variability. For all the subunits, the ICD showed much larger diversity than other regions. First, the length of this domain varied greatly in the subunits, with α5 the shortest and α4 the longest. Second, for this domain, the ω values for segments bordering TM3 or TM4 were relatively small, but they increased rapidly toward the center of the domain. For subunits α5, β2 and β3, the ICD was short (less than 100 aa) and had only a single peak locating in the middle of the domain. While for other subunits, the domain was not only much longer, but also had more complex patterns of ω value, usually with multiple peaks distributed over a large segment. The diversity of the patterns of ω value in this domain probably implicated a relatively relaxed functional or structural selective constraint in this region. Compared with other subunits, α7 had a unique distribution pattern of ω value, which was not only smaller, but also more uniformly distributed along the sequence. Additionally, subunits α4 and β2 had uniform and small ω values along the sequences except the ICD, indicating the presence of gentler and smaller selective pressure in the subunits but the ICD.

**Figure 1.**
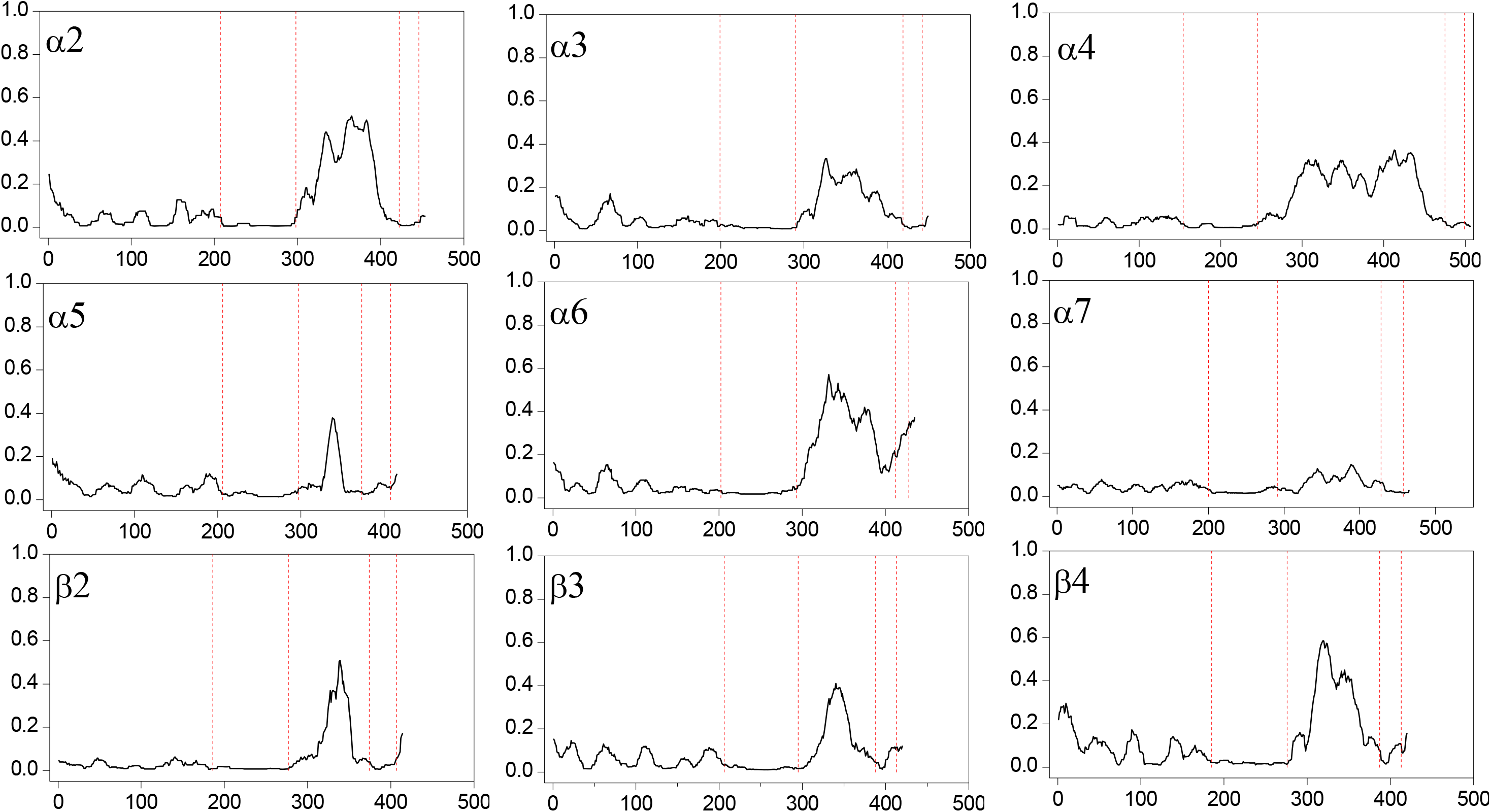
The ω values for each site along the sequence of a subunit across the species. A window-based analysis of ω values was performed to used to measure the local variation of selection pressure, with the window size defined as a sequence segment of 15-codon on the alignment of the same type of subunit. The vertical line shows the *f* values, and the horizontal line shows the amino acid site numbers of aligned sequences. Black and dashed arrows indicate the sequence regions which show the lower *f* values.

When each subunit was compared with other subunits, no sign of positive selection was detected except for subunit α5 and β2, on which the positively selected sites were found in the N-terminus and MX-M4 domain, respectively (Supplementary Table S2).

### 3.2. Potential function-specific sites in neuronal nAChR subunits

The current nAChR subunits are believed to be descended from a common ancestor, and the functional differentiation of these subunits arises from adaptive evolution after gene replication events. To further analyze the functional specificity in each subunit, it is necessary to identify sites potentially undergoing positive selection among those dominated by purifying selection. Earlier studies have showed that a nAChR subunit is more similar to the same type of subunit across species than to other types of subunit within the same species, which means each type of nAChR subunit from different species constitutes a separate branch on phylogenetic tree [12, 16]. Similar pattern could be seen for neuronal nAChR subunits (Supplemental Figure 3). Based on the phylogenetic features and experimental data available, the neuronal subunits could be grouped into five tribes, i.e., subunit α7, subunit β2 and β4, subunit α5 and β3, subunit α3 and α6, as well as α2 and α4 subunits.

In this study, the amino acids potentially related to evolution adaption and function differentiation in each subunit were identified via the branch-site model, a method able to detect the amino acid sites subject to positive selection along the sequence and among lineages. Briefly, based on the phylogenetic tree, each branch (i.e., nAChR subunit) was set to the foreground branch, with all the other branches being the background branch. On subunit α2, α3, α5, α6, β2 and β3, positive selection was detected for a number of sites, while for subunit α4, α7 and β2, none of the sites were found to be subject positive selection (Table 1). Most of the identified positively selected sites had a posterior probability between 0.5 and 0.95 in the BEB analysis (Supplementary Table S3).

**Table 1.** Positive selection detection in each subunit of vertebrate neural nAChR gene family using the branch-site model

| Foreground branch <sup>a</sup> | lnL | lnL (MA1) | LRT | p-value <sup>b</sup> | Positively selected sites <sup>c</sup> |
| --- | --- | --- | --- | --- | --- |
| α2 | -76122.79 | -76127.16 | 8.74 | 0.0031 | 100, 112, 115, 127, 141, 144, 147, 171, 185, 190, 205, 215, 131, 348, 351 |
| α3 | -76123.39 | -76127.34 | 7.90 | 0.0049 | 31, 101, 143*, 157, 205 |
| α4 | -76124.60 | -76125.92 | 2.64 | 0.10 | NA |
| α5 | -76107.62 | -76114.94 | 14.64 | 0.00013 | 28, 40, 91, 111, 132, 167, 182, 183, 186, 189, 196, 250, 265, 306, 322, 325, 617, 618 |
| α6 | -76122.78 | -76127.49 | 9.42 | 0.0021 | 35, 40, 143, 157*, 171, 346 |
| α7 | -76113.05 | -76113.37 | 0.64 | 0.43 | NA |
| β2 | -76123.62 | -76124.90 | 2.56 | 0.11 | NA |
| β3 | -76100.01 | -76108.20 | 16.38 | $5.20 \times 10^{-5}$ | 32, 34, 35, 64*, 115, 122, 183, 220, 221, 234, 273, 308, 333, 341, 342, 613, 615 |
| β4 | -76116.78 | -76120.99 | 8.42 | 0.0037 | 35, 53*, 58, 62*, 106, 132, 190, 214, 215, 218**, 235*, 594 |

Note. <sup>a</sup> In the first column, each subunit is selected as the foreground branch and the remaining subunits are background branches.
<sup>b</sup> A *p*-value < 0.05 means the subunit includes sites under positive selection.
<sup>c</sup> Sites with a BEB posterior probability > 0.95 are labeled with \*, and those higher than 0.99 are labeled with \*\*.
NA represents no positive selection detected. Residues are numbered according to their locations within human α4 subunit, as determined in the X-ray crystallographic structure of human α4β2 nicotinic receptor deposited in Protein Data Bank (PDB) (https://www.rcsb.org/structure/5KXI; PDB ID: 5KXI).

The positively selected sites in these branches could be largely grouped into two clusters. One cluster (Site 28, 31, 32, 34, 35, and 40; residues were numbered according to their locations within human α4 subunit, as specified in the structure of human α4β2 nAChR deposited in PDB, ID: 5KXI) gathered in the N-terminal α-helix. Another cluster (Site 182, 183, 186, 187, 189, and 190) flocked round the β8 strand neighboring the loop B and the loop F, particularly in the α5 branches. There were several scattered sites in the TMDs in the α5 and β3 branches, but not other branches. One site (N265) of α5 resided in the M1-M2 linker, negatively charged residues (D or E) in other subunits. E189 of α5 and E190 of α2 located at the first and second site of the β8-β9 loop, which was suggested to carry the allosteric site for nAChR potentiation by Ca^2+^ [41]. The location of D111 of α5 was homology to D111 of α9 subunit conferring high calcium permeability of mammalian α9α10 nAChRs [42].

Thus, these sites of α5 may be related to the function specificity of α5* receptors. E143 of α3, P157 of α6, K64 of β3 and S53, Q62, Q218 and K235 of β4 were under positive selection with posterior probabilities higher than 0.95, as calculated with the BEB approach.

### 3.3. Positive selection in neuronal nAChR subunit tribes

As specified above, the neuronal subunits could be grouped into five tribes. We further analyzed the functional specialization of these tribes and explored the molecular features responsible for the assortment of nAChR subunits via the branch-site model by detecting changes of ω ratios among sites in each tribe. In this step, the four tribes, i.e. α2α4, α3α6, α5β3, and β2β4, were set as the foreground branch and the remaining subunits were set as the background branch, respectively. Positive selection was detected in all branches (Table 2; Supplementary Table S4).

**Table 2.** Positive selection detection in each subunit of Vertebrate neural nAChR Gene Family using the branch-site model methodXS

| Foreground branch <sup>a</sup> | lnL | lnL (MA1) | LRT | P-value <sup>b</sup> | Positively selected sites <sup>c</sup> |
| --- | --- | --- | --- | --- | --- |
| $\alpha 2\alpha 4$ | -76094.21 | -76109.01 | 29.6 | $5.29 \times 10^{-8}$ | 28**, 47**, 56**, 102*, 129**, 137*, 139*, 188**, 210**, 213**, 215*, 582** |
| $\alpha 3\alpha 6$ | -76110.75 | -76115.05 | 8.60 | 0.0034 | 27*, 45*, 46*, 74**, 79**, 82*, 86*, 87**, 104*, 125**, 129**, 131*, 152*, 212*, 214*, 216*, 219*, 233**, 290*, 327*, 329*, 585*, 616* |
| $\alpha 5\beta 3$ | -76108.05 | -76110.79 | 5.48 | 0.019 | 54*, 112*, 244*, 251*, 266**, 288*, 296*, 335*, 592*, 612* |
| $\beta 2\beta 4$ | -76129.63 | -76120.55 | 18.16 | $2.03 \times 10^{-5}$ | 34*, 70*, 75**, 94*, 139*, 190*, 209*, 221*, 226*, 231**, 350* |

Note. <sup>a</sup> In the first column, each subunit tribe is selected as the foreground branch and the remaining subunits are background branches.
<sup>b</sup> A $p$ -value < 0.05 means the subunit includes sites under positive selection.
<sup>c</sup> Sites with a BEB posterior probability > 0.95 are labeled with \*, and those higher than 0.99 are labeled with \*\*.
NA represents no positive selection detected. Residues are numbered according to their locations within human $\alpha 4$ subunit, as determined in the X-ray crystallographic structure of human $\alpha 4\beta 2$ nicotinic receptor deposited in Protein Data Bank (PDB) (https://www.rcsb.org/structure/5KXI; PDB ID: 5KXI).

Of the twelve positively selected sites detected on α2α4 branch, most (28, 47, 56, 102,129,137, 139, 188, 210, 213 and 215) were in the N-terminal ECD, with only one site (582) located in TM4 domain. Twenty-three positively sites were identified for the α3α6 branch, while most sites were also found in N-terminal ECD, four sites (290, 327, 329 and 585) flocked together in the TMDs or ICD. In the α5β3 branch, ten positively sites were detected, with eight sites residing in the TMDs or TMD linker and only two sites (54 and 112) were in the ECD. Of the eleven positively selected sites detected in the β2β4 branch, ten sites were included in the ECD, and only one site (350) in TM4 domain. A site (70) located in the end of the β1 strand, which is the proved Ca^2+^ allosteric site of nAChRs. The dominant functional differentiation zone of β3 and α5 subunits was the TMDs. These sites may be bound up with α5 subunit and β3 subunit as the structural moiety, or they may be involved in the function regulation of the receptors including them.

Overall, most of these positively selected sites resided outside the pore of the receptor, with a large portion of sites were in the ECD (Figure 2A). In this domain, the β9 strand is conjectured as a region responsible for the functional differentiation and is involved in regulating functional specificity of receptors formed by α2, α4, α3 or α6 subunit (Figure 2B). Meanwhile, the β2 and β5 strand are regions that distinguished the α2α4 tribe from α3α6 tribe, as well as the α2α4 tribe from other tribes, respectively (Figure 2C); the β4-β5 linker distinguished the α2α4 tribe from the others (Figure 2D). A few sites reside at the TMDs, nearly all of which belonged to the α5β3 tribe, indirectly confirming the conservation and importance of the TMDs (Figure 2E). Five sites (74, 75, 190, 290 and 296) resided in the coupling region (Figure 2D).

**Figure 2.**
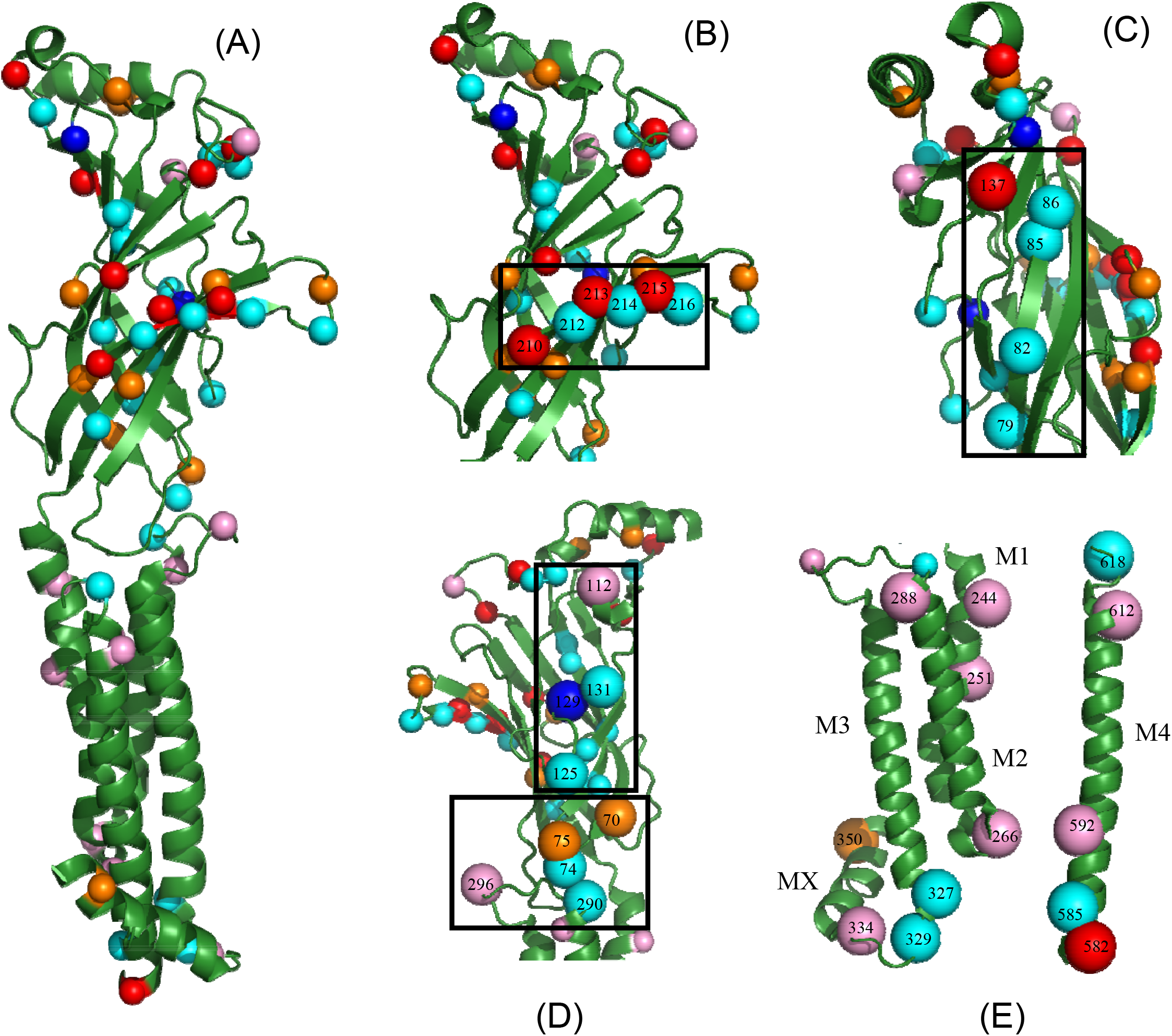
Mapping positively selected sites of the nAChR subunit onto the 3D molecular structure. The subunit is composed of an amino-terminal extracellular domain, four transmembrane domains (i.e., TM1-TM4), an intracellular domain and a short extracellular carboxyl-terminus. The extracellular domain contains eleven β-strands (termed as β1-β10 and β6’ respectively). Residues within each tribe are shown as colored spheres, with those in the α2α4 tribe shown as red, the α3α6 tribe as cyan, the β2β4 tribe as orange and the α5β3 tribe as pink. (A) Distribution of residues on the subunit; (B) residues of the β9 strand; (C) residues of β2 and β5 strand(loop E) of the coupling region; (D) residues of the β4-β5 linker (upper) and the coupling region (lower); (E) residues of the TMDs.

## 4. Discussion

The diversity in function and the combinations of nAChR subunits are responsible for the highly variable kinetic, electrophysiological and pharmacological properties of the receptors. Since the current neuronal subunits in vertebrates were descended from common ancestors, comparing them in the context of evolution could help us understand the specificity of each subunit. By analyzing the sequence variations in the neuronal nAChR subunits, we revealed the common and unique features existed in the subunits and identified a profile of sites may be related to their molecular structure and function.

### 4.1. Evolution of neuronal nAChRs subunits in vertebrates

Our results indicate that the intracellular domain of the subunits may be in the process of adaptation. Of the neuronal subunits, α7, α4 and β2 is constrained by strong selective pressure, which results in rather small variation along the sequence except the intracellular domain. Some sites of TM4 of α6, β3 and β4, especially α6 subunit, showed functional or structural relaxed selective constraints. It is known that there is a strong link between channel gating and lipid-protein interaction. Considering the interaction between TM4 of the subunits, such features may be related to the channel gating of the receptor. Evidences of positive selection after vertebrate radiation have been found in the α5 and β2 sequences using the site models. The two identified regions, the N-terminus and the MX-M4, suggest the existence of adaptive process of these subtypes in the evolution of vertebrates. Actually, subunit α5 is found to be closely connected with the evolution of the brain [43]. Thus, the increase in complexity of the cholinergic systems should be considered as part of evolution of the nervous system.

### 4.2. Molecular evolution of neuronal nAChR subunits diversity

One the most compelling evidence of subunit functional specificity comes from the fact that sequences of nAChR family members are more similar to the same type of subunit across species than different subunits within the same species. As we know, the nAChR subunits were generated by gene duplication and the multiple types of neuronal nAChR subunit have been conserved through evolution of vertebrate. It is likely that the diversity of neuronal nAChRs subunits is the consequence of adaptive evolution to different patterns of selection pressure imposed to each subunit. In our study, the signatures of positive selection related to subunit differentiation were captured by the branch-site analyses. For subunit α2, α3, α5, α6, β3 and β4, a number of sites subject to positive selection were identified. However, for subunit α4, α7 and β2, no site under significant positive selection was detected. Actually, subunit α7 may be an ancestral type subunit [44], and nicotinic receptors formed by subunit α4 and β2 are the most abundant subtype in the brain and α4 and β2 could represent a ‘fossil’ expression, which was present in most areas of the ancestral brain [12]. These subunits may have been under strong functional or structural selective constraints in vertebrates. Consistent with such site model results, subunit α4, β2 and α7 are in the strongly adaptive selective pressure during vertebrate, as indicated by the relatively small and uniformly distributed ω values along the sequence except the ICD, and the small ω value of α7 along its whole sequence.

Analysis at residue level revealed several positively selected sites in each subunit. The short segment 180-183 (loop B) of α7-α4 chimeras was shown to alter the pharmacology of all agonists, independent of their chemical structure [45]. The contribution of the region deserves to underscore the role of positive selection of S182 of α5, M183 of α5 and β3. E190 of α7 is the Ca^2+^ allosteric site for potentiating the function of α7 nAChRs, and the β8-β9 loop is the Ca^2+^ allosteric region for nAChRs potentiation [41]. D111 of α9 confers high calcium permeability of mammalian α9α10 nAChRs [42]. It deserves that the effect of E189 of α5, E190 of α2 and D111of α5 on their formed receptor.

A site cluster locates at N-terminal α-helix. One allosteric site is located near of the N-terminal helix of the extracellular domain on the α7 receptors. Ligand binding at this site causes a conformational change of the α-helix as the fragment wedges between the α-helix and a loop homologous to the main immunogenic region of the muscle α1 subunit [46], which illustrates the place may have role in allosteric modulation or in the functional features favoring subtype differentiation.

### 4.3. Functional differentiation in neuronal nAChRs subunit tribes

Several whole genome duplications occurred at the beginning of vertebrate evolution, which probably were the origin of the initial diversity in many gene families [47–48]. The phylogenetic tree in tribes of the neuronal nAChR subunits are congruent with the groups of subunits, defined on the basis of biochemical, functional or pharmacological data. Evidences of positive selection have been found in all tribes before vertebrate radiation. These decide on functional differentiation of these subunits, thus the proper of formed receptor.

The 211-219 segment of α7-α4 chimera contributes to the relative ACh and nicotine affinities. The 209–223 segment accounted for approximately half the difference in toxin sensitivity between α2* and α3* nAChRs, in particular, K213 and I216 of α3 are critical in determination of sensitivity to toxin blockade[49]. A site P210 in the β9 strand of the outer β-sheet, allows for rapid desensitization of receptors, for example α7 nAChRs [50]. The proof of positive selection of T/V210, Y212 and I216 of α3α6 tribe, Y213, H214 and S/T215of α2α4 tribe indicates that the β9 strand of the α subunit of heteromeric receptors regulates pharmacological diversity and specificity and the β9 strand of subunits is in favor of desensitization kinetics through residues on the β9 strand that does not contribution directly to ligand binding. The β9 strand may be valuable for the function of nAChRs.

Meanwhile, residues of the vestibule β4-β5 loop interact with passing Ca^2+^ in the mammalian α9α10 nAChRs [42], and the vestibule of α7 receptors exist a negative allosteric site and corresponds to a previously identified site involved in positive allosteric modulation of the bacterial homolog ELIC [46], justifying the positive selection on the β4-β5 loop of the α3α6 tribe has important role in the formed receptors, such as Ca^2+^ permeability or the allosteric modulation. At the β2-α4 interface in (α4) _2_ (β2) _3_ receptors, H137 of α4 is a NS9283-enhanced allosteric site [51], which is located at the beginning of loop-E (β5 strand). It is unexpected that some positively selected sites exist on the β2 strand (loop-D) of α3α6 tribe and the β5 strand (loop-E) of α2α4 tribe. They certainly connect with the property of formed receptors, such as diverse assembly, like α6β3α4 and α6β3α4β2 receptors.

Experiments have shown that α5 and β3 subunits contribute to channel characteristic of nAChRs [52]. However, the functional differentiation region of α5β3 tribe also lies in the TMDs. It is indicative of the TMDs of α5 and β3 subunits play a potential role in α5* and β3* receptors. Each subunit has two acidic amino acids in the M1-M2 linker, where are important to determine ion selectivity and conductance, but the β3subunit has three. Mutation E268A of α7 abolishes the permeability of a7 receptors to Ca^2+^, but preserves that to monovalent cations [44]. E266 of the α5β3 tribe maybe contribute to the relative ion current through the pore.

The event of positive selection also occurs in the coupling region. The coupling region is termed as the bridge of the signal transduction, participating in the diverse function of receptors [52]. This residue D70 belongs to a Ca^2+^ allosteric site, which may explain potentiation by Ca^2+^ in α7 receptors much stronger than in (α4)_2_(β2)_3_ [53]. The importance of the region is qualified to consider the role of positive selection.

Positive selection is in favor of subtype-specific functions. The combined action of gene duplication and positive selection may have originated the actual vertebrate subtypes, which are conserved by strong purifying selection.

Undeniably, there are several limitations in this study. First, only twelve vertebrate species were analyzed in the current work. Given that the genomes of more and more species are available, nAChR subunits from more species should be analyzed. As the identification genes encoding nAChR subunits is an ongoing process, results from the current study should be treated in the same way. Second, in our study, the subunits in other species were selected based on their sequence homology to the corresponding human subunits, we could not rule out the possibility that some subunits may have function different than their counterparts in human. Especially, due to genome duplication, some fish species have much larger nAChR gene family than other species [54]. Third, although some results from this study are consistent with those from other techniques such as molecular biology and genetic analysis, they should be further verified by other approaches in the future.

## 4.4. Conclusions

In summary, in this study, we revealed the common and unique features between the members of neuronal nAChRs gene family. We have demonstrated the existence of purifying selection in all the neuronal nAChR subunits in vertebrate, but the functional constraints are not uniform within a subunit, and the subunits are subject to different patterns of selection pressures in evolution. Positive selection seems to act on specific positions favoring special characteristic of each subtype and each tribe, which may contribute to the diversity and specificity of the pharmacological properties, desensitization kinetics, and other peroperties of the subunits and receptors formed by them. The coupling region contributes to the signal transduction and its positive selection seems to bear on functional differentiation of subunits prior to the radiation of vertebrate orders. In the process of evolution of vertebrates, α7, α4 and β2 subunits possess a strong adaptability. This study may provide useful information for understanding the structure and function, as we all as evolution of nAChRs, and could bring in new insights for the design of new drugs targeting on nAChRs.

## Acknowledgements

This study was supported in part by grants from National Key Research and Development Program of China (No.2016YFC0906300), National Natural Science Foundation of China (No. 31271411 & 91746205). The funding bodies played no role in the design of the study and collection, analysis and interpretation of data, or in writing of the manuscript.

