## Supplementary figures and images for "Analysis of the Functional Differentiation Sites in Vertebrate Neuronal Nicotinic Acetylcholine Receptor Subunits"

### Supplemental Figures

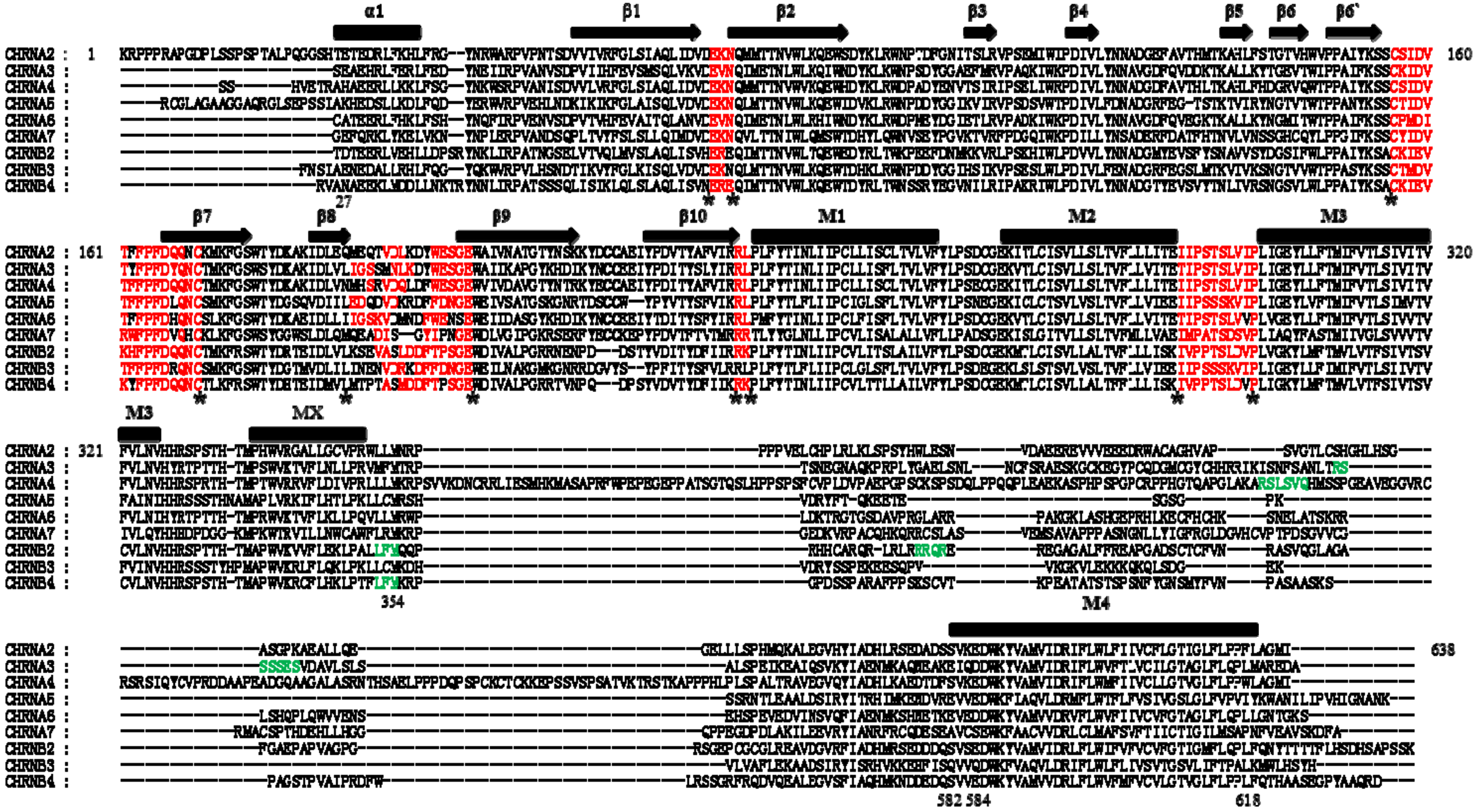

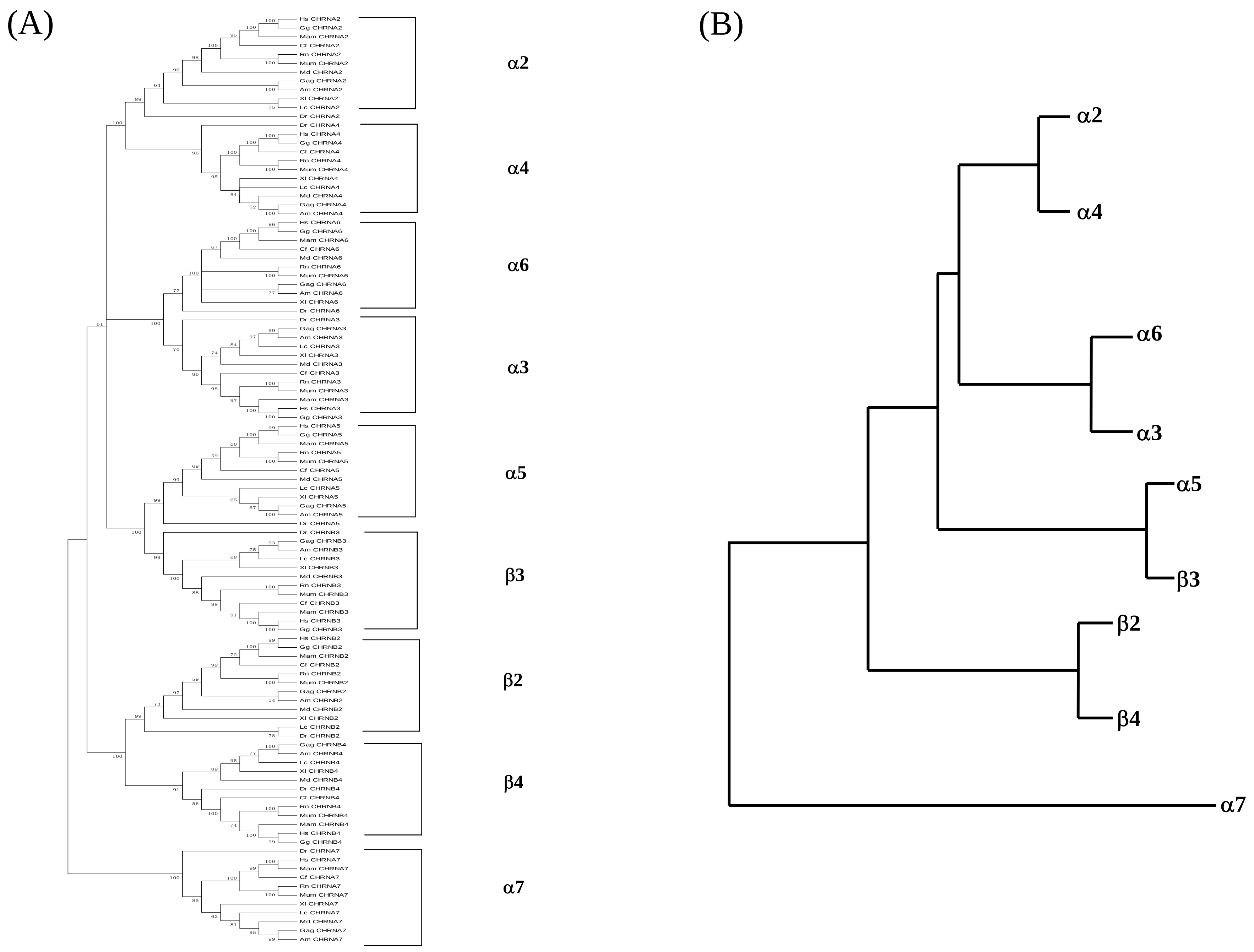


Figure S2

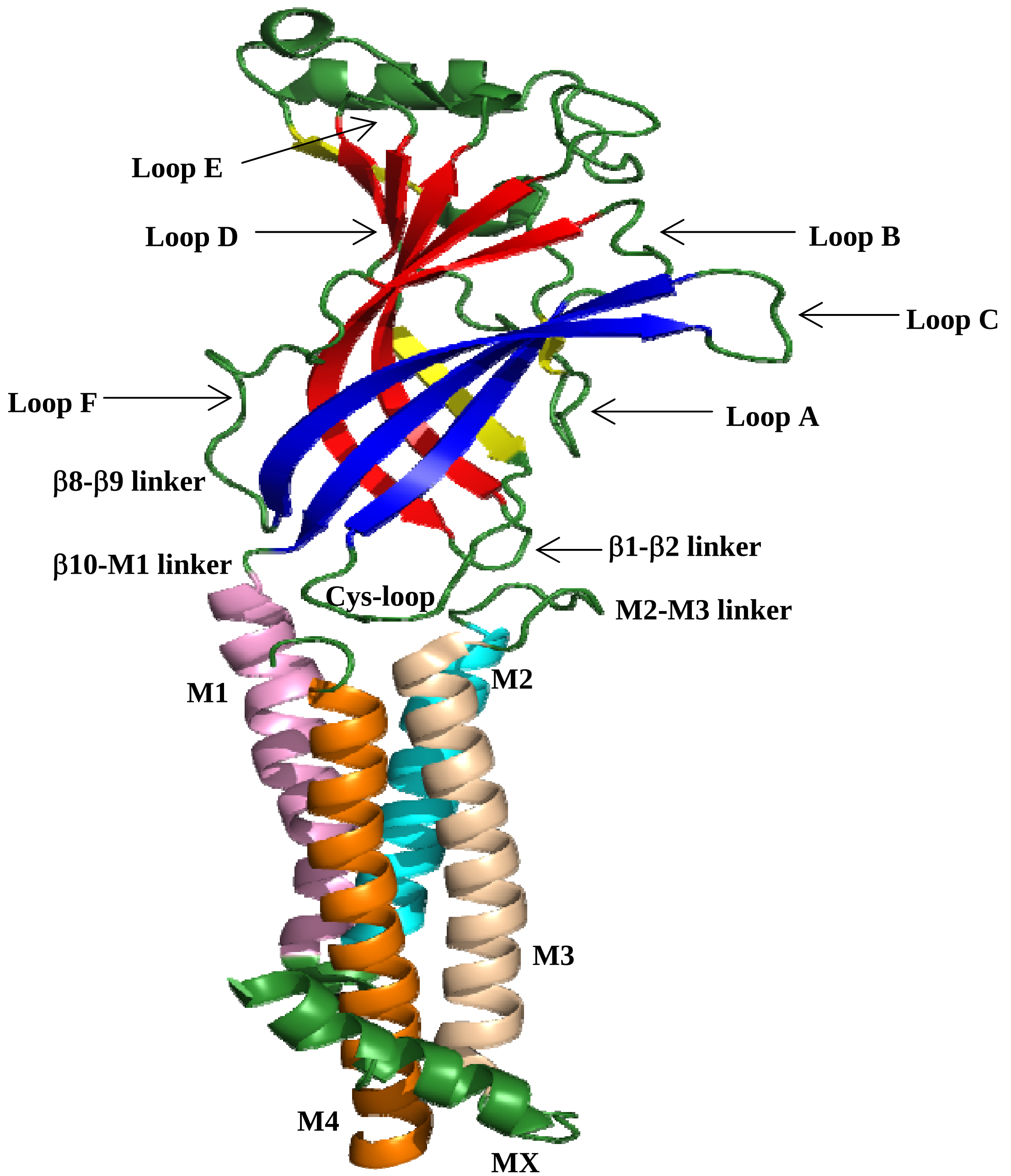
