## Supplemental Tables for "Analysis of the Functional Differentiation Sites in Vertebrate Neuronal Nicotinic Acetylcholine Receptor Subunits"

Table S1. Sequences of nine nicotinic acetylcholine receptor subunits from twelve representative vertebrates

| Subunit | Species | Sequences of the subunits* |
|---|---|---|
| CHRNA2 | Homo sapiens | NM_000742 |
| | Gorilla gorilla | XM_004046819 |
| | Macaca mulatta | CM001260 |
| | Rattus norvegicus | NM_133420 |
| | Mus musculu | NM_144803 |
| | Canis lupus familiaris | XM_005635889 |
| | Monodelphis domestica | XM_001380021 |
| | Gallus gallus | NM_204815 |
| | Alligator mississippiensis | XM_006274660 |
| | Xenopus laevis | XM_018264125 |
| | Latimeria chalumnae | XM_006010203 |
| | Danio rerio | NM_001040327 |
| CHRNA3 | Homo sapiens | NM_000743 |
| | Gorilla gorilla | XM_004056608 |
| | Macaca mulatta | XM_015142832 |
| | Rattus norvegicus | NM_052805 |
| | Mus musculus | NM_145129 |
| | Canis lupus familiaris | XM_014118840 |
| | Monodelphis domestica | XM_007478003 |
| | Gallus gallus | NM_204416 |
| | Alligator mississippiensis | XM_006261354 |
| | Xenopus laevis | XM_018251424 |
| | Latimeria chalumnae | XM_006001841 |
| | Danio rerio | XM_001921279 |
| CHRNA4 | Homo sapiens | NM_000744 |
| | Gorilla gorilla | XM_004062519 |
| | Macaca mulatta | −− |
| | Rattus norvegicus | NM_024354 |
| | Mus musculus | NM_015730 |
| | Canis lupus familiaris | XM_014107318 |
| | Monodelphis domestica | XM_001376166 |
| | Gallus gallus | NM_204814 |
| | Alligator mississippiensis | XM_019484414 |
| | Xenopus laevis | XM_018237044 |
| | Latimeria chalumnae | XM_005993470 |
| | Danio rerio | NM_001048063 |
| CHRNA5 | Homo sapiens | NM_000745 |
| | Gorilla gorilla | XM_004056611 |
| | Macaca mulatta | XM_015142833 |
| | Rattus norvegicus | NM_017078 |
| | Mus musculus | NM_176844 |

| | | |
|---|---|---|
| | Canis lupus familiaris | XM_005628321 |
| | Monodelphis domestica | XM_007478206 |
| | Gallus gallus | NM_204415 |
| | Alligator mississippiensis | XM_006261353 |
| | Xenopus laevis | NM_001093080 |
| | Latimeria chalumnae | XM_006001846 |
| | Danio rerio | NM_001017885 |
| | Homo sapiens | NM_004198 |
| | Gorilla gorilla | XM_004046970 |
| | Macaca mulatta | XM_001099152 |
| | Rattus norvegicus | NM_057184 |
| | Mus musculus | NM_021369 |
| CHRNA6 | Canis lupus familiaris | XM_539951 |
| | Monodelphis domestica | XM_001381900 |
| | Gallus gallus | NM_205364 |
| | Alligator mississippiensis | XM_006276967 |
| | Xenopus laevis | XM_018252934 |
| | Latimeria chalumnae | —— |
| | Danio rerio | NM_001042684 |
| | Homo sapiens | NM_000746 |
| | Gorilla gorilla | —— |
| | Macaca mulatta | NM_001032883 |
| | Rattus norvegicus | NM_012832 |
| | Mus musculus | NM_007390 |
| CHRNA7 | Canis lupus familiaris | XM_545813 |
| | Monodelphis domestica | XM_001377478 |
| | Gallus gallus | NM_204181 |
| | Alligator mississippiensis | XM_014599894 |
| | Xenopus laevis | XM_018253016 |
| | Latimeria chalumnae | XM_006011688 |
| | Danio rerio | NM_201219 |
| | Homo sapiens | NM_000748 |
| | Gorilla gorilla | XM_004026831 |
| | Macaca mulatta | XM_015109918 |
| | Rattus norvegicus | NM_019297 |
| | Mus musculus | NM_009602 |
| CHRNB2 | Canis lupus familiaris | XM_547565 |
| | Monodelphis domestica | XM_001373116 |
| | Gallus gallus | NM_204813 |
| | Alligator mississippiensis | AKHW03005445 |
| | Xenopus laevis | NM_001094830 |
| | Latimeria chalumnae | XM_005996497 |
| | Danio rerio | XM_005169754 |
| CHRNB3 | Homo sapiens | NM_000749 |

| | | |
|---|---|---|
| | Gorilla gorilla | XM_004046969 |
| | Macaca mulatta | XM_015145273 |
| | Rattus norvegicus | NM_133597 |
| | Mus musculus | NM_173212 |
| | Canis lupus familiaris | XM_539952 |
| | Monodelphis domestica | XM_007476412 |
| | Gallus gallus | NM_204812 |
| | Alligator mississippiensis | XM_006276963 |
| | Xenopus laevis | XM_018243476 |
| | Latimeria chalumnae | XM_006007586 |
| | Danio rerio | NM_201220 |
| CHRNB4 | Homo sapiens | NM_000750 |
| | Gorilla gorilla | XM_004056612 |
| | Macaca mulatta | XM_015142823 |
| | Rattus norvegicus | NM_052806 |
| | Mus musculus | NM_148944 |
| | Canis lupus familiaris | XM_847455 |
| | Monodelphis domestica | XM_007477997 |
| | Gallus gallus | NM_204819 |
| | Alligator mississippiensis | XM_014605947 |
| | Xenopus laevis | XM_018252852 |
| | Latimeria chalumnae | XM_006001842 |
| | Danio rerio | XM_691901 |

* For subunit $\alpha 4$ of *Macaca mulatta*, $\alpha 6$ of *Latimeria chalumnae*, and $\alpha 7$ of *Gorilla gorilla*, the corresponding nucleotide sequence is either unavailable yet or is excluded for low quality.

Table S2. Positively Selected Sites Detected in Each Subunit of Vertebrate neural nAChR Gene Family

| Subunits | M8 Model | | M7 Model | | LRT | P-value | Domain of positively selected sites |
|---|---|---|---|---|---|---|---|
| | lnL | Parameter | lnL | Parameter | | | |
| α2 | -9553.85 | p0=1.00 p=0.14 q=0.89 (p1=0.00) w = 1.00 | -9553.85 | p=0.14 q = 0.88 | 0.00 | 1.00 | NA |
| α3 | -8667.41 | p0=0.99 p=0.19 q=2.43 (p1 = 0.01) w = 1.00 | -8668.29 | p=0.18 q = 1.98 | 1.76 | 0.42 | NA |
| α4 | -11437.68 | p0=1.00 p=0.19 q= 1.39 (p1 = 0.00) w = 1.00 | -11437.68 | p=0.19 q = 1.39 | 0.00 | 1.00 | NA |
| α5 | -7997.93 | p0=0.95 p=0.22 q= 2.44 (p1= 0.050) w = 4.14 | -8028.49 | p=0.16 q= 0.99 | 61.12 | $5.35\times10^{-14}$ | N- terminus |
| α6 | -8471.71 | p0= 0.97 p=0.25 q=1.86 (p1 = 0.030) w = 1.00 | -8472.62 | p=0.22 q = 1.33 | 1.82 | 0.40 | NA |
| α7 | -7646.65 | p0=1.00 p =0.26 q= 5.07 (p1 = 0.00) w = 1.016 | -7646.65 | p=0.26 q = 5.07 | 0.00 | 1.00 | NA |
| β2 | -8177.47 | p0 = 0.96 p =0.17 q = 3.61 (p1 = 0.040) w = 1.00 | -8183.74 | p=0.13 q= 1.54 | 12.54 | 0.0019 | MX-M4 |
| β3 | -7971.53 | p0=0.99 p=0.19 q= 2.00 (p1 = 0.010) w = 1.22 | -7972.12 | p=0.19 q = 1.75 | 1.18 | 0.56 | NA |
| β4 | -9607.86 | p0=0.97 p=0.23 q= 1.70 (p1 = 0.030) w = 1.19 | -9610.06 | p=0.20 q= 1.16 | 4.40 | 0.11 | NA |

Table S3. Positively Selected Sites Detected in Each Subunit of Vertebrate neural nAChR Gene Family using the branch-site model method

| Foreground branch | Positively Selected Sites | Posteriori probability | Location on the molecular |
|---|---|---|---|
| | 100 | 0.510 | β2-β3 linker |
| | 112 | 0.512 | β3-helix |
| | 115 | 0.538 | β3-β4 linker |
| | 127 | 0.681 | β4-β5 linker |
| | 138 | 0.599 | β5-β6 linker |
| | 141 | 0.587 | β6 strand |
| | 147 | 0.828 | β6-β6` linker |
| α2 | 171 | 0.583 | β7 strand |
| | 185 | 0.927 | β8 strand |
| | 190 | 0.799 | β8-β9 linker |
| | 205 | 0.552 | β9 strand |
| | 215 | 0.502 | β9 strand |
| | 131 | 0.561 | β10 strand |
| | 348 | 0.579 | MX |
| | 351 | 0.934 | MX |
| | 31 | 0.833 | N-terminal α-helix |
| α3 | 101 | 0.55 | β2-β3 linker |
| | 143 | 0.973 | β6 strand |
| | 157 | 0.857 | β6`-β7 linker |
| | 35 | 0.866 | N-terminal α-helix |
| | 40 | 0.523 | α1-β1 linker |
| α6 | 143 | 0.514 | β6 strand |
| | 157 | 0.983 | β6`-β7 linker |
| | 171 | 0.644 | β7 strand |
| | 346 | 0.576 | MX |
| α5 | 28 | 0.809 | N-terminal α-helix |
| | 40 | 0.822 | α1-β1 linker |
| | 91 | 0.894 | β2-β3 linker |
| | 111 | 0.768 | β3-helix |
| | 132 | 0.831 | β4-β5 linker |
| | 167 | 0.712 | β7 strand |
| | 182 | 0.709 | β7-β8 linker |
| | 183 | 0.872 | β7-β8 linker |
| | 186 | 0.864 | β8 strand |
| | 189 | 0.741 | β8-β9 linker |

| | | | |
|---|---|---|---|
| | 196 | 0.843 | β8-β9 linker |
| | 250 | 0.807 | M1 |
| | 265 | 0.786 | M1-M2 |
| | 306 | 0.793 | M3 |
| | 322 | 0.778 | M3 |
| | 325 | 0.772 | M3 |
| | 617 | 0.808 | C-terminal |
| | 618 | 0.883 | C-terminal |
| β3 | 32 | 0.939 | N-terminal α-helix |
| | 34 | 0.846 | N-terminal α-helix |
| | 35 | 0.871 | N-terminal α-helix |
| | 64 | 0.963 | β1 strand |
| | 115 | 0.817 | β3-β4 linker |
| | 122 | 0.870 | β4-β5 linker |
| | 183 | 0.915 | β7-β8 linker |
| | 220 | 0.863 | β9-β10 linker |
| | 221 | 0.846 | β9-β10 linker |
| | 234 | 0.898 | β10 strand |
| | 273 | 0.950 | M2 |
| | 308 | 0.860 | M3 |
| | 333 | 0.852 | M3-MX |
| | 341 | 0.863 | MX |
| | 342 | 0.822 | MX |
| | 613 | 0.874 | M4 |
| | 615 | 0.849 | C-terminal |
| β4 | 35 | 0.75 | N-terminal α-helix |
| | 53 | 0.981 | α1-β1 linker |
| | 58 | 0.543 | β1 strand |
| | 62 | 0.958 | β1 strand |
| | 106 | 0.52 | β3 strand |
| | 132 | 0.512 | β4-β5 linker |
| | 190 | 0.912 | β8-β9 linker |
| | 214 | 0.734 | β9 strand |
| | 215 | 0.535 | β9 strand |
| | 218 | 0.994 | β9-β10 linker |
| | 235 | 0.974 | β10 strand |
| | 594 | 0.53 | M4 |

Table S4. Positively Selected Sites Detected in Each Tribe of vertebrate neural nAChR gene family using the branch-site model method

| Foreground branch | Positively Selected Sites | Posteriori probability | space distribution |
|---|---|---|---|
| α2α4 | 28 | 0.995 | N-terminal α-helix |
| | 47 | 0.994 | α1-β1 linker |
| | 56 | 0.96 | α1-β1 linker |
| | 102 | 0.999 | β2-β3 linker |
| | 129 | 0.991 | β4-β5 linker |
| | 137 | 0.981 | β5 strand |
| | 139 | 0.952 | β5 strand |
| | 188 | 0.997 | β8 strand |
| | 210 | 0.998 | β9 strand |
| | 213 | 0.998 | β9 strand |
| | 215 | 0.979 | β9 strand |
| | 582 | 0.993 | M4 |
| α3α6 | 27 | 0.990 | N-terminal α-helix |
| | 45 | 0.960 | α1-β1 linker |
| | 46 | 0.982 | α1-β1 linker |
| | 74 | 0.998 | β1-β2 linker |
| | 79 | 0.999 | β2 strand |
| | 82 | 0.984 | β2 strand |
| | 86 | 0.982 | β2 strand |
| | 87 | 0.999 | β2 strand |
| | 104 | 0.982 | β2-β3 linker |
| | 125 | 0.991 | β4-β5 linker |
| | 129 | 0.992 | β4-β5 linker |
| | 131 | 0.986 | β4-β5 linker |
| | 152 | 0.980 | β6` strand |
| | 212 | 0.980 | β9 strand |
| | 214 | 0.973 | β9 strand |
| | 216 | 0.979 | β9 strand |
| | 219 | 0.973 | β9-β10 linker |
| | 233 | 0.998 | β10 strand |
| | 290 | 0.973 | TM2-TM3 linker |
| | 327 | 0.973 | TM3 |
| | 329 | 0.991 | M3-MX linker |
| | 585 | 0.968 | TM4 |
| | 616 | 0.959 | C-terminal |
| α5β3 | 54 | 0.981 | α1-β1 linker |
| | 112 | 0.989 | β3-β4 linker |

| | | | |
|---|---|---|---|
| | 244 | 0.966 | TM1 |
| | 251 | 0.990 | TM1 |
| | 266 | 0.996 | TM1-TM2 |
| | 288 | 0.968 | TM2 |
| | 296 | 0.986 | TM2-TM3 linker |
| | 335 | 0.954 | M3-MX linker |
| | 592 | 0.966 | TM4 |
| | 612 | 0.958 | TM4 |
| | 34 | 0.960 | N-terminal α-helix |
| | 70 | 0.983 | β1 strand |
| | 75 | 0.991 | β1-β2 linker |
| | 94 | 0.985 | β2-β3 linker |
| | 139 | 0.982 | β5 strand |
| β2β4 | 190 | 0.958 | β8-β9 linker |
| | 209 | 0.973 | β9 strand |
| | 221 | 0.989 | β9-β10 linker |
| | 226 | 0.998 | β10 strand |
| | 231 | 0.991 | β10 strand |
| | 350 | 0.965 | MX |
| | 34 | 0.960 | N-terminal α-helix |

**Figure Legends:**

**Supplemental Figure S1.** Alignment of the neuronal nAChR subunit sequences. The alignment result reveals several highly conserved features. The transmembrane domains are quite conservative with non-polar residues (e.g., Phe, Val, Ile and Leu) occupying a large proportion. The coupling region is also conserved across all the subunits. Among the five segments of the coupling region, four (i.e., β1-β2 linker, Cys-loop, β10-M1 linker, and M2-M3 linker) are conserved not only in all subunits of every species, but also in each subunit across all species analyzed. For the β8-β9 linker, although several sites showed larger variability, most residues were also conserved; especially, the last five sites were conserved in all subunits across species. Several motifs are conserved in the large intracellular domain. The ER export motif LMF was detected in sequences of subunit α2, α4, β2 and β4. While this motif was presented in subunit β4 of all species, it was absent in subunit β2 of rodent, α2 of mammals and α4 of primates. A motif related to ER retention and/or retrieval (RRQR) was found in subunit β2 of mammals and primates. A 14-3-3 binding motif (RSSSSES) and a similar RSLSVQ motif were conserved in subunit α3 and α4 of non-amphibians, respectively. These motifs may be involved in regulating nAChRs maturation in vertebrate species.

**Supplemental Figure S2.** Phylogenetic tree of neuronal nAChR subunits. (A). Subunits in all the species analyzed are shown; (B). The collapsed phylogenetic relationships among vertebrate neuronal nAChR subunits, in which the branches corresponding to the same subunits of different species are condensed up to one node.

**Supplemental Figure S3.** The framework composes of an amino-terminal extracellular domain, four transmembrane domains (termed as TM1-TM4), a large and variable intracellular domain between the TM3 and TM4 domains, as well as a short extracellular carboxyl-terminus. The extracellular domain contains eleven β-strands (termed as β1-β10 and β6′ respectively) following an N-terminal α-helix. The region containing acetylcholine binding sites of nAChRs, also termed as the ligand binding domain (LBD), is located at the interface between two neighboring subunits and contributed by three loops (termed Loop A, B and C) from the principal subunit and three loops (termed Loop D, E and F) from the complementary subunit. The interacting regions between LBD and the transmembrane domain are named as ‘coupling region’, which is composed of five segments, i.e., β1-β2 linker, Cys-loop (formed by the linker between strand β6′ and β7, as well as part of strand β7), β8-β9 linker, β10-TM1 linker, and the TM2-TM3 linker. These linkers are involved in relaying the effect of ligand binding to the ion channel of the receptor. The TM2 helices of the five subunits in a receptor are closely packed to pave the inner circle of the ion channel pore, while TM1, TM3 and TM4 concentrically form the outer circle. The intracellular domain, together with the coupling region, may impart features specific to each subunit.
